# Separation of heterogeneous *Aggregatibacter actinomycetemcomitans* outer membrane vesicles reveals that toxin sorting is driven by surface-associated DNA

**DOI:** 10.1101/2022.07.11.499574

**Authors:** Justin B. Nice, Shannon M. Collins, Samuel M. J. Agro, Anxhela Sinani, Spencer D. Moros, Leah M. Pasch, Angela C. Brown

## Abstract

*Aggregatibacter actinomycetemcomitans* is a Gram-negative oral bacterium associated with localized aggressive periodontitis as well as some systemic diseases. The strains of *A. actinomycetemcomitans* most closely associated with periodontal disease tend to produce more of a secreted leukotoxin (LtxA) than isolates from healthy carriers, suggesting a key role for this toxin in disease progression. LtxA is secreted via a type 1 secretion system, from the bacterial cytosol across both the inner and outer membranes in a single step into the supernatant, where it is able to interact with host cells. Upon secretion, some of the toxin associates with the bacterial cell membrane, enabling its release in association with the surface of outer membrane vesicles (OMVs), small, spherical vesicles derived from the outer membrane. We have previously observed that the highly leukotoxic *A. actinomycetemcomitans* strain JP2 produces two populations of OMVs, a highly abundant population of small (<100 nm in diameter) OMVs, and a less abundant population of large (>300 nm in diameter) OMVs. Here, we have investigated the association of LtxA with the two populations of OMVs varying in size. Our results indicate that surface-associated DNA drives the selective sorting of LtxA to large OMVs.

## 1. Introduction

The Gram-negative bacterium, *Aggregatibacter actinomycetemcomitans*, is associated with aggressive forms of periodontitis, particularly in adolescents [1-4], as well as systemic infections including endocarditis [5]. This organism produces several virulence factors, including a leukotoxin (LtxA), which selectively kills white blood cells [6, 7], thus inhibiting an effective host immune response. The amount of LtxA that is produced by different strains of *A. actinomycetemcomitans* has been correlated to the strains’ association with periodontal disease [2, 4, 8-12], suggesting an important function of this virulence factor in pathogenesis, especially in juveniles.

Like other members of the repeats-in-toxin (RTX) family of proteins [13], LtxA is secreted via a type 1 secretion system (T1SS) across both the inner and outer membranes of the bacterium in a single step [14, 15]. In this secreted form, the toxin interacts with host cell membranes through recognition of both cholesterol [16, 17] and an integrin receptor, lymphocyte function associated antigen-1 (LFA-1) [18-20].

In addition to this “free” form of LtxA, the toxin has been observed to be delivered to host cells through association with outer membrane vesicles (OMVs) [21, 22]. We and others have found that LtxA is located on the surface of the OMVs [21, 23], consistent with prior reports that LtxA has an affinity for the bacterial cell surface, reported to be driven by interactions with the lipopolysaccharide (LPS) [11] and surface-associated DNA [23, 24], in addition to environmental conditions, such as pH [14] and iron concentration (FeCl_3_) [25]. Thus, it seems that after secretion via the T1SS, some of the toxin associates with the bacterial cell surface, localizing it to the OMV surface following vesicle formation.

Importantly, the OMVs released by *A. actinomycetemcomitans* are enriched in LtxA relative to the concentration on the outer membrane (OM) [21, 26], suggesting some type of sorting mechanism to promote OMV association. Several reports have found that although OMVs are derived from the bacterial OM, significant differences in protein and lipid composition between the OMV and OM can exist, indicating that certain molecules localize to the OMV, while others remain on the OM. For example, the LPS structure present in the OMVs produced by both *Porphyromonas gingivalis* and *Pseudomonas aeruginosa* differs from that of the parent cell [27, 28]. In both cases, OMVs were enriched in negatively charged species of LPS relative to the parent cell. Likewise, OMVs produced by *A. actinomycetemcomitans* strain JP2, contained three unidentified lipids that were not present in the bacterial OM sample [26]. In addition, certain proteins have been reported to selectively sort into OMVs [27, 29, 30].

In addition to compositional differences between the OM and OMVs, several groups have reported that OMVs released by a single organism can exhibit heterogeneous physical properties, particularly size [31-34]. Rather than a continuous distribution of sizes, this heterogeneity has manifested in bimodal size distributions. Interestingly, *Helicobacter pylori* was observed to produce a heterogeneous population of OMVs, with diameters ranging from 20 to 450 nm; separation of these OMVs by density gradient centrifugation (DGC) demonstrated that the smaller OMVs had a less diverse protein content than larger OMVs, and the two populations of OMVs entered host cells via different mechanisms [31]. In our previous work, we noted that the JP2 strain of *A. actinomycetemcomitans* likewise releases heterogeneous OMVs with a bimodal size distribution consisting of a minority population with diameters of approximately 325 nm and a majority population with diameters of approximately 100 nm [21]. Here, we sought to separate these OMV populations with the goal of determining whether LtxA associates with one or both populations and identifying the factors driving this association.

Previous studies have employed DGC to separate OMV populations by density (and therefore, size) [31, 32, 34]. However, this method is time-consuming and often results in incomplete separation of vesicles [31, 33]. More recently, size exclusion chromatography (SEC) has been utilized to remove impurities that pellet with vesicles during ultracentrifugation [35-37]. In this study, we used an ultrafine SEC resin to separate OMV subpopulations by size [38] so that we could explore the mechanism driving the selective packaging of LtxA into subpopulations of OMVs. This technique enabled us to efficiently and reproducibly separate the two OMV subpopulations to demonstrate an increased association with LtxA with the larger subpopulation of OMVs. In addition, we demonstrated that this sorting is regulated by variations in the concentration of surface-associated DNA between the OMV subpopulations.

## 2. Materials and Methods

### 2.1 Bacterial Culture

*A. actinomycetemcomitans* strains JP2 (serotype b), ATCC 33384 (serotype c) [39], and AA1704, an isogenic *ltxA* mutant [40] of the JP2 strain, were grown under identical conditions. The growth media consisted of 30 g/L trypticase soy broth (BD Biosciences, Franklin Lakes, NJ, USA) with 6 g/L yeast extract (BD Biosciences), 0.4% sodium bicarbonate (VWR, Radnor, PA, USA), 0.8% dextrose (BD Biosciences), 5 μg/mL vancomycin (Sigma-Aldrich, St. Louis, MO, USA), and 75 μg/mL bacitracin (Sigma-Aldrich). Cultures were first grown in a candle jar for 16 hr at 37 °C, then transferred at a ratio of 3:50 to fresh media and grown an additional 24 hr at 37 °C outside of the candle jar. A 1.5 L culture was grown to the late exponential phase (optical density at 600 nm, OD_600_ of 0.68-0.72).

### 2.2 OMV Purification

To purify OMVs from the culture supernatant, the bacteria were pelleted by centrifugation twice at 10,000 x *g* at 4 °C for 10 min. The supernatant was next filtered through a 0.45 μm filter. The bacteria-free supernatant was concentrated using Amicon® 50 kDa filters (MilliporeSigma, Burlington, MA) and then ultracentrifuged at 105,000 x *g* at 4 °C for 30 min. Pellets were pooled in phosphate-buffered saline (PBS, pH 7.4), ultracentrifuged again, and resuspended in 1 mL of PBS.

### 2.3 Dynamic Light Scattering (DLS)

An ALV/CGS-3 goniometer system was used to determine the diameter distributions of the OMVs. Samples were suspended in PBS and measured for three minutes at a wavelength of 632.8 nm and a scattering angle of 90°. Size distributions were calculated on the ALV software (ALV-5000/E, ALV-GmbH, Langen, Germany, 2001), using a number-weighted regularized fit with the coated sphere assumption (membrane thickness r^*^=5 nm) [41, 42]. The number-weighted fit corrects for the fact that larger particles scatter more light than smaller particles by normalizing the particles by count rather than scattering.

### 2.4 Size Exclusion Chromatography

SEC was used to separate OMVs by size; a 1.5 cm x 50 cm (bed volume 85 mL) was packed with Sephacryl™ S-1000 superfine resin (GE Healthcare, Chicago, IL, USA) [43] and equilibrated with two bed volumes of PBS. A 2-mL OMV sample was loaded, eluted with PBS, and one-mL fractions were collected. Fractions were analyzed for lipid, LtxA, and surface-associated DNA concentrations, as described below.

### 2.5 Lipid Content

The percentage of lipid in each SEC fraction was measured using the FM 4-64™ dye (ThermoFisher Scientific, Waltham, MA). First, 50 μL of each fraction was incubated with FM 4-64™ (0.1 mg/mL) for 15 sec. Following incubation, the fluorescence of the sample was measured on a Tecan plate reader with an excitation wavelength of 515 nm and an emission wavelength of 640 nm. The fluorescence intensity of each fraction was divided by the summed intensities to calculate a percentage of total lipid in each fraction.

### 2.6 Surface-Associated DNA Content

The amount of DNA on the surface of the OMV fractions was measured using TOTO™-1 Iodide (ThermoFisher Scientific). A TOTO™-1 stock solution was made by mixing 1 μL TOTO™-1 (1 mM in DMSO) with 250 μL Tris-EDTA buffer (TE buffer, 0.1 M Tris, 0.1 M EDTA, pH 8.0). The stock solution was mixed in equal volume ratios with the OMV fractions and incubated for 10 min at room temperature. Following incubation, the fluorescence was measured using a Quantamaster® 400 spectrofluorometer (PTI Horiba, Edison, NJ) with an excitation wavelength of 514 nm and an emission wavelength of 533 nm.

### 2.7 Protein Characterization

The LtxA concentration in each SEC fraction was measured using an enzyme-linked immunosorbent assay (ELISA). Fractions were incubated in a MaxiSorp Immuno 96-well plate (ThermoFisher Scientific) for 3 hr, washed five times with ELISA wash buffer (25 M Tris, 150 mM sodium chloride, 0.1% fatty-acid free bovine serum albumin (BSA)) then blocked in 1% BSA in the same buffer. The plate was then incubated with an anti-LtxA antibody [44] in 1% BSA/buffer overnight at 4 °C. Following five washes with ELISA wash buffer, the plate was incubated in goat anti-mouse horseradish peroxidase (GAM-HRP) at a 1:5000 ratio (SouthernBiotech, Birmingham, AL). Lastly, the plate was imaged using 1-Step™ Ultra TMB ELISA substrate solution (ThermoFisher Scientific) until signal appeared, then the reaction was stopped using 2 M sulfuric acid. The absorbance at 450 nm was measured on a Tecan plate reader. The resulting absorbance of each fraction was divided by the summed absorbances to calculate a percentage of total LtxA in each fraction.

LtxA content was also analyzed by Western blotting. Sodium dodecyl sulfate polyacrylamide gel electrophoresis (SDS-PAGE) was performed using 7.5% acrylamide gels. Western blotting for LtxA was accomplished by transferring the proteins to a nitrocellulose membrane overnight. The blots were washed three times in tris-buffered saline with 0.1% tween (TBST) and then blocked with blotto solution (5% dried milk in TBST) for 1 hr. LtxA was then detected using a monoclonal anti-LtxA antibody [44] overnight at 4°C, followed by GAM-HRP for 1 hr. The blot was imaged using SuperSignal™ West Dura substrate (ThermoFisher). To measure the amount of outer membrane protein-A (OmpA) in each fraction, a similar procedure was used, with the anti-Gram-negative OmpA antibody (111228, Antibody Research Corporation, St. Charles, MO) at a final concentration of 0.24 μg/mL, followed by goat anti-rabbit horseradish peroxidase (GAR-HRP, SouthernBiotech). Densitometry analysis on the blots was accomplished using ImageJ [45].

### 2.8 LtxA Binding Experiments

To determine the role of LPS in LtxA association with OMVs, the OMVs were treated with polymyxin B at a concentration of 4 mg/mL at 37°C for 1 hr [11] before separation by SEC, as described above. To determine the role of DNA in LtxA association with OMVs, the OMVs were treated with DNase I (Sigma-Aldrich) at a final concentration of 100 U/mL at 37°C for 1 hr [23]. Following treatment, OMVs were separated by SEC as described above.

### 2.9 Statistical Analysis

Statistical analysis was performed with Sigma-Plot, using an unpaired two-tailed student’s *t*-test, where p values ≤0.01 were considered to be statistically significant.

## 3. Results

### 3.1 SEC Separation of OMVs Yields Two Populations

OMVs were collected from the end of the exponential phase of growth, purified by ultracentrifugation, and separated using the Sephacryl S-1000 SF packed column [43]. One-mL fractions were collected, and the presence of OMVs in each fraction was determined using the FM™ 4-64 dye. Fig. 1A shows the FM™ 4-64 fluorescence values of each fraction. Two peaks were observed, consistent with OMVs of two distinct sizes. The size distribution of the OMVs found in each fraction was then measured by DLS. The number-weighted DLS distributions of each fraction are shown in Fig. 1B, where cool colors represent earlier fractions and warm colors represent later fractions. DLS analysis confirmed that the fractions eluting at 34-36 mL column volume contained mostly large OMVs (>250 nm) while fractions eluting at 44-52 mL were comprised of small OMVs (<250 nm). Together, these results indicate that SEC was effective in separating small and large OMVs.

**Figure 1.**
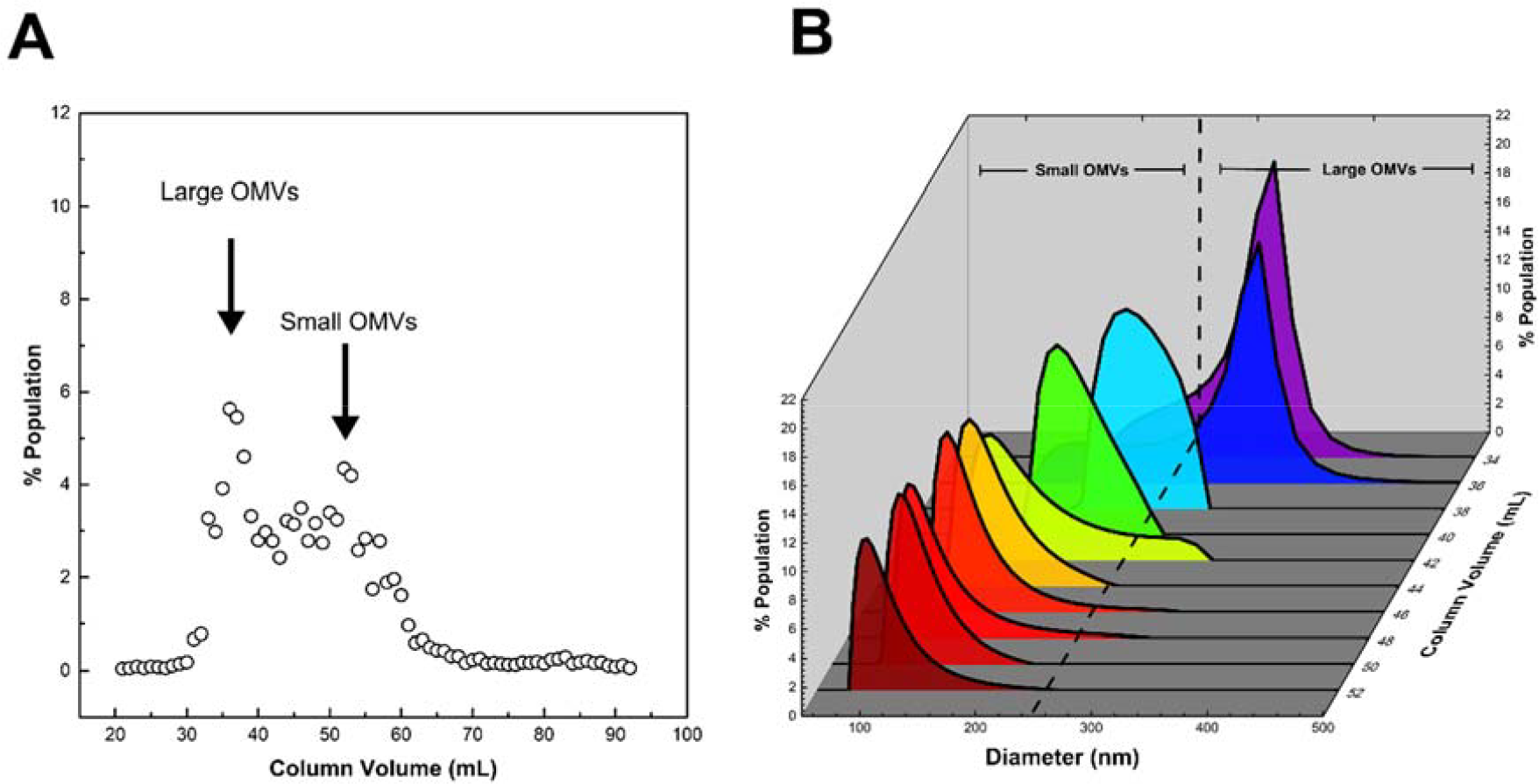
Separation of *A. actinomycetemcomitans* OMVs by SEC and analysis of OMV subpopulation size by DLS. (A) Fractional lipid compositions of each SEC fraction. The lipid concentrations in each fraction were determined using FM™ 4-64 dye fluorescence intensity. The intensity of each fraction was compared to the summed intensity to calculate a percentage of total lipid in each fraction. The data shown are representative results from n=3 trials. (B) Number-weighted DLS size distributions of each SEC fraction. Each plot shows the percentage of OMVs in each fraction with a given diameter. Cool colors represent earlier fractions; warm colors represent later fractions. The data shown are representative results from n=3 trials.

### 3.2 Large OMVs Are Enriched in Full-length LtxA

The LtxA concentration in each vesicle fraction was compared using ELISA with an anti-LtxA antibody [44]. The total ELISA signal was normalized to 100%, and the distribution of LtxA by fraction is shown in Fig. 2A, along with the lipid compositions. Most of the LtxA eluted at 34-36 mL, fractions which contained mostly large OMVs, with a very small amount of LtxA co-eluting with small OMVs in column volumes 44-52 mL. These results demonstrate that LtxA associates primarily with the large OMVs.

**Figure 2.**
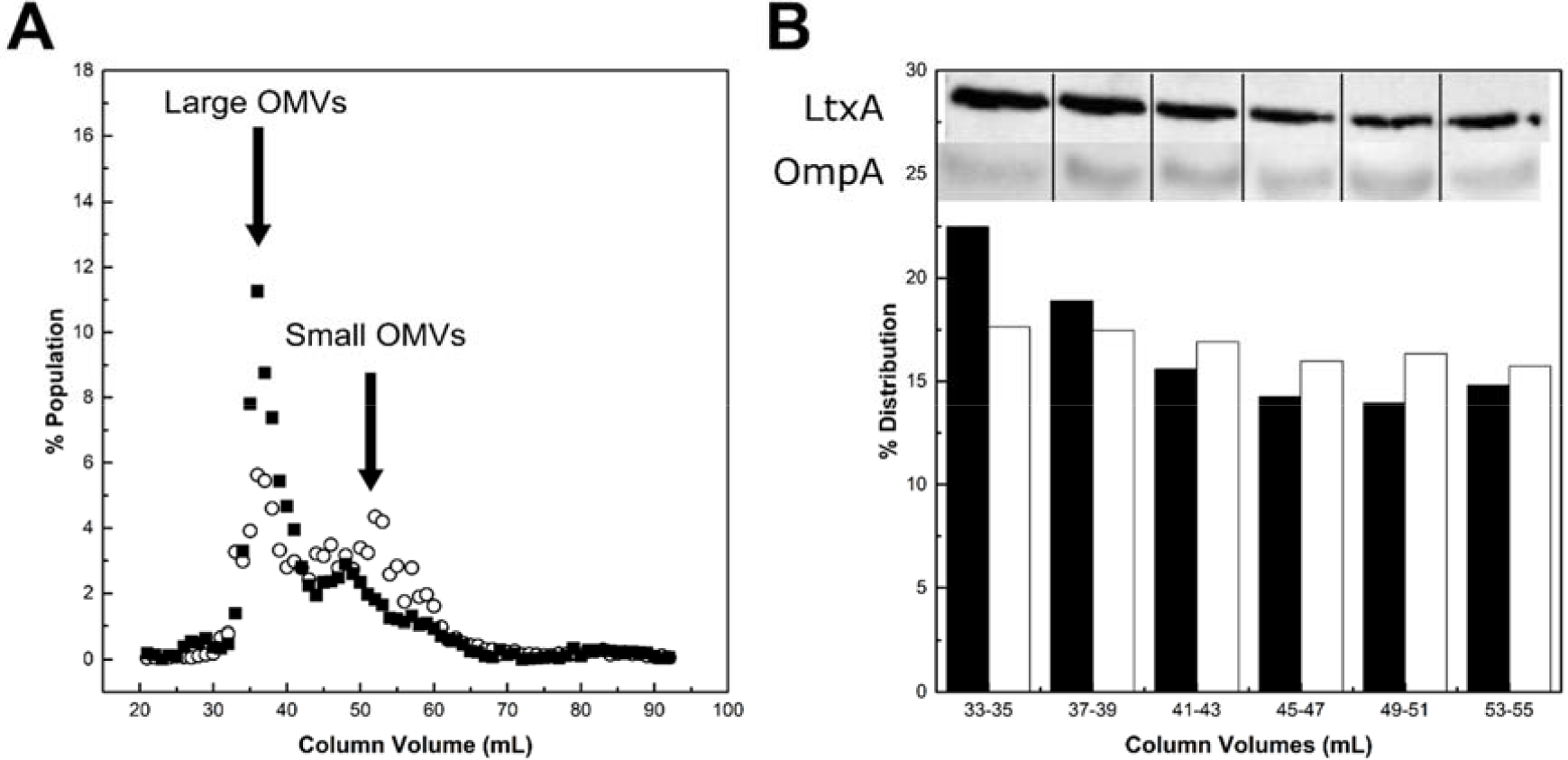
Protein composition of SEC fractions. (A) LtxA. The concentration of LtxA in each fraction was determined by ELISA (black squares). The intensity of each fraction was compared to the summed intensity to calculate a percentage of the total LtxA concentration in each fraction. The data shown are representative results from n=3 trials. (B) LtxA and OmpA. Insets: Western blots of lipid-normalized combined fractions for either LtxA or OmpA. Densitometry analysis of the LtxA (black bars) and OmpA (white bars) Western blot bands. The data shown are representative results from n=3 trials.

It is possible that the increased amount of LtxA detected in the large OMV fractions could simply be due to the fact that the larger OMVs have ten times more surface area than the smaller OMVs, and thus more binding area. We therefore performed a Western blot on combined fractions, where each sample was normalized to have the same lipid concentration, based on the FM™ 4-64 dye intensity. This normalization allowed us to look at the amount of LtxA per lipid in each fraction, thus eliminating the differences in surface area. The signal strength was quantified using densitometry, and the total signal of all lanes was normalized to 100% (Fig. 2B). The Western blot demonstrated that those fractions containing primarily large OMVs (33-35 mL) were enriched with full length LtxA (114 kDa), while those comprising small OMVs (41-51 mL) contained much less LtxA (Fig. 2B, inset). In parallel, a Western blot for outer membrane protein A (OmpA) was performed on the same fractions; OmpA is found on both the bacterial OM and OMVs, and is often used as an OMV marker [34, 46]. The OmpA concentration was the same in all samples, demonstrating that, unlike LtxA, OmpA was sorted to the OMVs uniformly and independently of vesicle size (Fig. 2B, inset).

### 3.3 Large OMVs Are Produced Independently of LtxA

LtxA is a membrane-active protein that is able to induce curvature in model membrane systems [47]. We therefore hypothesized that the preferential association of LtxA with large OMVs might be due to LtxA on the surface of the bacterial cells mediating large OMV formation through curvature induction. To test this hypothesis, we compared JP2 OMVs with those produced by AA1704, a non-LtxA expressing isogenic mutant of JP2 [40], as well as *A. actinomycetemcomitans* strain 33384, a minimally leukotoxic, serotype c strain [39]. JP2 and AA1704 differ only in the expression of LtxA, while JP2 and 33384 differ in both LtxA expression and LPS structure. DLS was used to determine the distribution of vesicle diameters (Fig. 3A). Fig. 3B represents concentrations of the small (<250 nm) and large (>250 nm) OMVs calculated from the DLS distributions. We found that although AA1704 does not produce LtxA, it was able to produce both large and small OMVs, similarly to JP2. In contrast, ATCC 33384 (serotype c) did not produce any large vesicles. These results demonstrate that, contrary to our hypothesis, LtxA is not directly involved with the production of large OMVs; rather the toxin must have a preferential affinity for the surface of large OMVs. We therefore next sought to understand what mediates this differential affinity.

**Figure 3.**
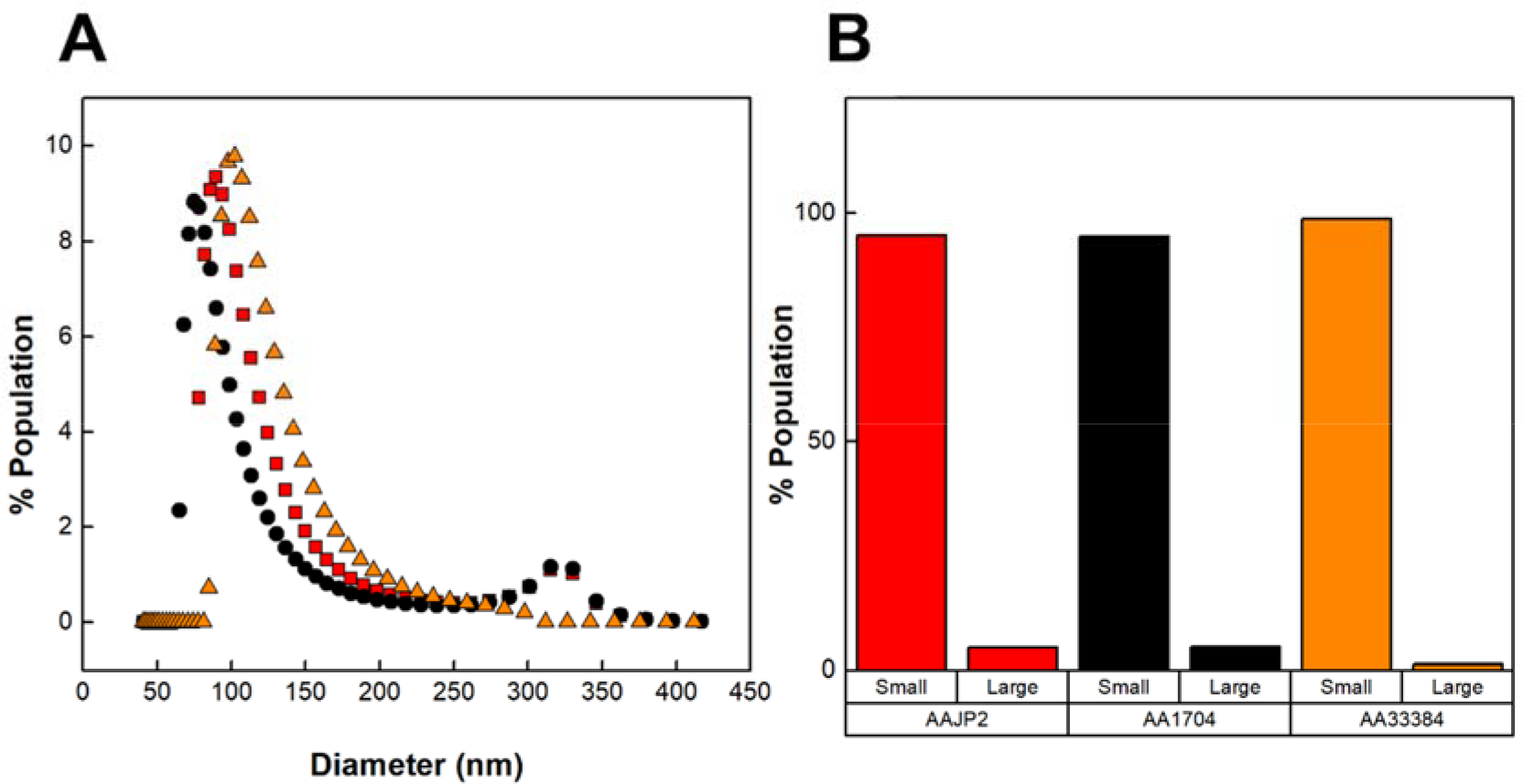
DLS distributions of OMVs produced by various strains of *A. actinomycetemcomitans*. **(A) Number-weighted DLS distributions of OMVs**. Each plot shows the percentage OMVs produced by JP2 (black circles), AA1704 (red squares), and ATCC 33384 (orange triangles) with a particular diameter. The data shown are representative results from n=3 trials. **(B) Concentrations of large and small OMV subpopulations**. Large and small concentrations were calculated from the DLS distributions. “Large” OMVs are defined as those larger than 250 nm, while “small” OMVs are defined as those smaller than 250 nm in diameter. The data shown are representative results from n=3 trials.

### 3.4 LtxA Binding to Large OMVs is Mediated by Surface-Associated DNA

We previously demonstrated that LtxA is associated with the surface of JP2 OMVs [21], a finding that is consistent with prior reports that polymyxin B and DNase I treatments removed LtxA from the surface of *A. actinomycetemcomitans* cells and vesicles [11, 23, 24]. Therefore, we investigated whether the selective sorting of LtxA to large OMVs is mediated by either LPS or surface-associated DNA. We tested both conditions by first pretreating JP2 OMVs with polymyxin B or DNase I and measuring the resulting changes in the LtxA elution profile by SEC.

Fig. 4A shows a representative SEC elution profile after polymyxin B-treatment of the OMVs. The lipid and LtxA concentrations of untreated OMVs are included for reference. Following polymyxin B-treatment, the LtxA concentration in the fractions comprising large OMVs decreased slightly, and the LtxA concentration in fractions 70-83 mL, corresponding to free LtxA (that is, unassociated with OMVs), increased slightly. This result indicates that polymyxin-mediated neutralization of LPS removed a small amount of LtxA from the surface of the large OMVs.

**Figure 4:**
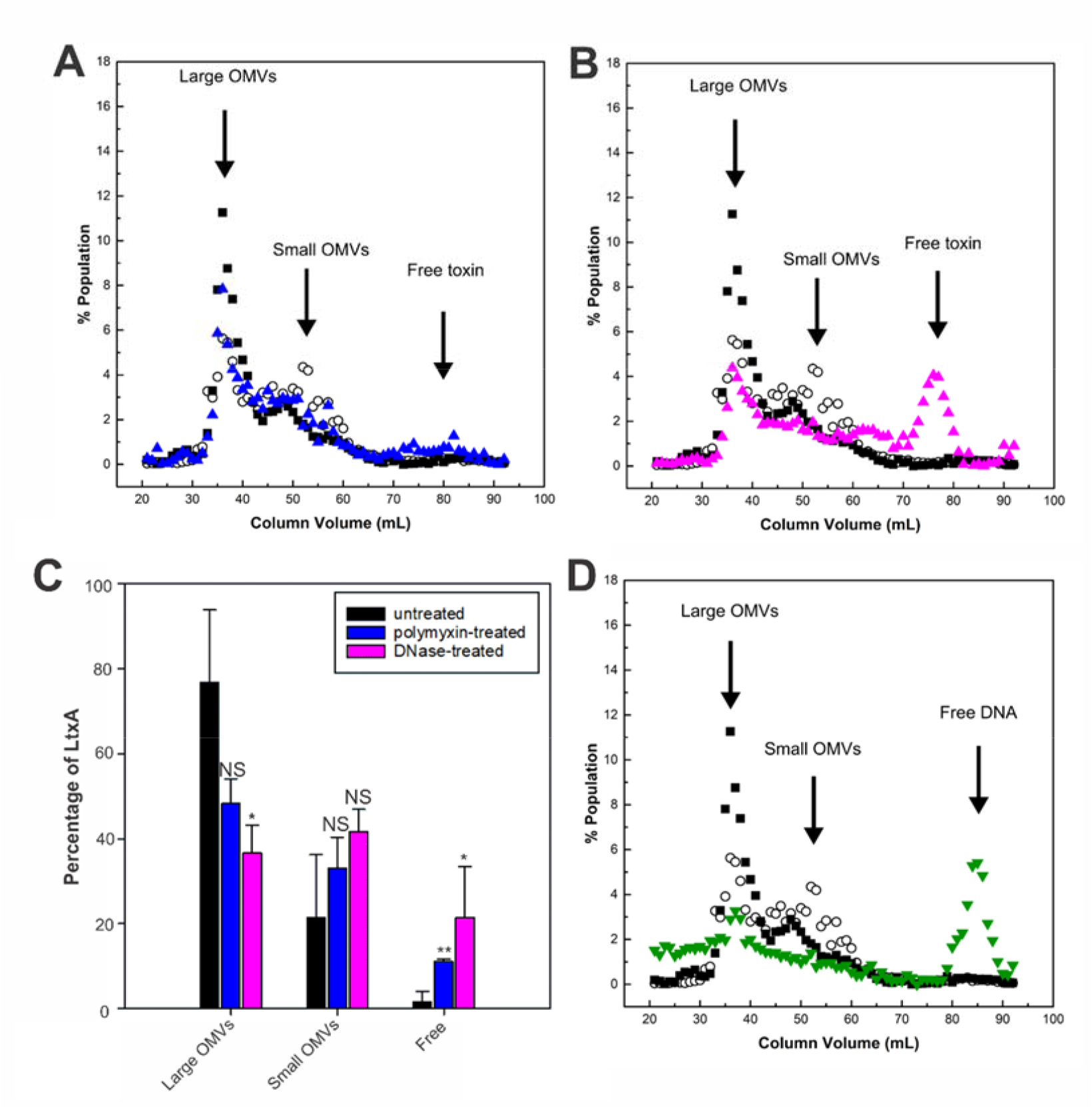
Mechanisms of LtxA binding to OMVs. (A) Polymyxin B-treated JP2 OMVs run on SEC. Polymyxin treated (blue triangles), untreated LtxA concentration (black squares), and lipid concentration (empty circles) are replotted for reference. The data shown are representative results from n=3 trials. (B) DNase I-treated JP2 OMVs run on SEC. DNase I treated (pink triangles). (C) Percentage of LtxA associated with large OMVs, associated with small OMVs, or unassociated before (black) or after polymyxin (blue) or DNase (pink) treatment. Each data point represents the mean ± standard deviation. *, p < 0.05; **, p < 0.01; NS, not significant, relative to respective untreated value. (D) DNA concentration on the surface of OMVs by TOTO-1. DNA concentration (green triangles). The data shown are representative results from n=3 trials.

Because polymyxin only removed a small amount of LtxA from the large OMVs, we next investigated the role of surface-associated DNA in the association of LtxA with large OMVs. The OMVs were treated with DNase I before separation by SEC to determine the effect of surface-associated DNA on the affinity of LtxA for OMVs. Fig. 4B also includes the lipid and LtxA concentrations of untreated OMVs for reference. After DNase I-treatment, the LtxA concentration in the fractions containing large OMVs decreased drastically, while the amount in the fractions containing free LtxA (70-83 mL) increased correspondingly. Fig. 4C shows the fraction of LtxA associated with large OMVs, associated with small OMVs, or unassociated (“free”) before and after polymyxin- or DNase-treatment for three separate trials. These results demonstrate that LtxA binding to the surface of the large vesicles is affected by the composition of the surface and is primarily DNA-driven.

The importance of surface-associated DNA was further assessed by measuring the amount of surface-associated DNA in each OMV fraction. Using the membrane impermeable nucleic acid stain, TOTO-1 [48], we found that DNA was concentrated in the fractions containing the large OMVs, but not in the fractions containing small OMVs (Fig. 4D). Additionally, we observed a significant amount of DNA elution in very late fractions, after free LtxA, which is likely unassociated DNA that co-purifies with the OMVs.

## 4. Discussion

OMVs have emerged as an important factor in bacterial virulence [46, 49, 50]. In particular, their role in delivery of toxins to host cells has been established as an important aspect of virulence. OMV-mediated transport of toxins enables delivery to the cytoplasm of the host cell [49] in a form that is protected from harsh extracellular conditions, including proteases [33, 46, 51]. In many cases, it has been observed that the delivery of OMV-associated toxin to host cells differs from that of free toxin [51-53]. For example, we have observed that OMV-associated LtxA is delivered to host cells in a cholesterol- and LFA-1 independent manner [21, 22], unlike free LtxA which requires both of these molecules [16, 19, 20]. While the mechanisms by which free toxins are delivered to host cells have been well established, the delivery of OMV-associated toxins to host cells is currently a subject of intense focus. Thus, understanding the nature of the association of toxins with OMVs is essential in elucidating the mechanisms by which these OMVs are produced and interact with host cells.

In this work, we found that LtxA is located predominately on the larger OMVs produced by *A. actinomycetemcomitans* strain JP2. This differential association is driven by variations in the amount of surface-associated DNA between the two OMV populations. A previous study of *A. actinomycetemcomitans*, strain D7SS, found that LtxA was associated with OMVs with reduced density, that is, larger OMVs, compared to another exotoxin, cytolethal distending toxin (Cdt) [34]. The D7SS strain is a nonfimbriated smooth colony mutant of a clinical isolate of *A. actinomycetemcomitans* [54], suggesting that heterogeneous sorting of LtxA on OMVs is a common and clinically relevant phenomenon. In addition, sorting of RTX toxins to certain OMVs may be conserved and related to their secretion mechanism, as it was reported that the RTX toxin, α-hemolysin, associates preferentially with large *E. coli* vesicles, along with some components of the T1SS machinery [33].

To investigate the process mediating the association of LtxA with the larger OMVs, we examined LPS and surface-associated DNA. It has been reported that LtxA can be extracted from the JP2 bacterial cells via polymyxin treatment [11], suggesting that LtxA association with LPS on the bacterial cell is mediated by interactions with LPS. However, we found that only a small amount of LtxA was removed from the OMVs by this treatment. We then investigated the role of OMV surface-associated DNA in this process, as DNA has also been reported to mediate the binding of LtxA to the bacterial cell membrane [23, 24]. In this case, we found that DNase treatment greatly reduced the amount of LtxA associated with the large OMVs, indicating that LtxA binds to DNA on the surface of the large OMVs. Consistent with this observation, we found that the large OMVs contain much more DNA on their surface than the smaller OMVs.

Current work is focused on understanding the biophysical and biochemical processes involved in DNA association with the large OMVs. We hypothesize that variations in the LPS structures and charge of the two OMV types regulate the binding of DNA. LPS variations between the OMVs and OM have been reported previously, for several organisms, including *P. gingivalis* [55], *E. coli* [56], and *P. aeruginosa* [28]. TLC analysis of *A. actinomycetemcomitans* JP2 OMVs revealed the presence of three lipids, which could include LPS, that were not present in the OM of these cells [46]. We suspect that similar variations in the LPS structures of the large and small OMVs, and the resulting variations in charge, might regulate DNA binding to the large OMVs. The origin of this DNA is also not currently known. Bitto *et al*. found that during the exponential phase of growth, DNA associates with OMVs [57]; they hypothesized that actively dividing bacteria shed DNA which then associates with OMVs. We are currently investigating variations in DNA and LtxA composition of the *A. actinomycetemcomitans* JP2 OMVs as a function of growth phase.

Recent work on OMVs has demonstrated the importance of studying subpopulations of vesicles rather than the bulk populations of OMVs produced during bacterial growth. The Kaparakis-Liaskos group found that separation of *H. pylori* OMVs by DGC yielded large and small vesicles, and they demonstrated that vesicle size affects both protein cargo and mechanism of entry into host cells [31, 32]. This group found that heterogeneity arises, at least in part, due to variations in the types of vesicles released during different growth phases, with more heterogeneous vesicles being released during early stages of growth, and less heterogeneous vesicles being released later in growth [32].

OMV heterogeneity and the resulting differences in cell uptake suggests that distinct populations of OMVs may serve different purposes. However, little work has been accomplished in this area due to the technical difficulties of completing such studies. One such difficulty is the efficient separation of a sufficient quantity of OMVs to complete the studies. DGC is commonly used to remove protein aggregates and flagellae [35], and it can also be used to separate OMV subpopulations [31, 34]. Drawbacks of DGC include that it is time-consuming, results in low yield [58], and requires optimization of fractionation protocols for each strain or species of bacteria [35]. Thus, optimization itself is time-consuming as each iterative change in the DGC protocol is arduous. This technique is also not well-suited for scale-up [58]. SEC has been proposed as a simpler and more efficient method for purifying extracellular vesicles (EVs) [59, 60]. Commercially available qEV columns (Izon Science Ltd, New Zealand) have been employed to purify EVs [61]; the use of these columns resulted a high-throughput method to obtain a pure population of EVs [62]. Recently, Singorenko, *et al*. compared qEV SEC columns with DGC for the purification of bacterial vesicles, and found that the columns were much quicker and better able to remove contamination [35]. The major drawback of the qEV columns, however, was the incomplete separation of a heterogeneous population into distinct subpopulations [35]. In our studies, we packed a longer column with Sephacryl™ S-1000 superfine resin in order to quickly and efficiently purify and separate OMV subpopulations [38]. The results we have reported here demonstrate that separation by SEC was rapid (about 4 hr), resulted in complete separation of OMV subpopulations, and removed free DNA contamination.

## 5. Conclusion

Using the SEC technique, we were able to separate two populations of *A. actinomycetemcomitans* JP2 OMVs efficiently and repeatably, enabling us to characterize sorting of LtxA between the vesicle populations. We anticipate that this approach will enable us to efficiently purify distinct populations of OMVs to better understand LtxA delivery to host cells via OMVs and to investigate the function and delivery processes of the different OMV populations.

## Acknowledgements

This work was supported by the National Institutes of Health grants DE025275 (ACB) and the National Science Foundation grant 1554417 (ACB). We would like to thank Lee Graham, Lehigh University Department of Biological Sciences Lab Manager for technical assistance.

## Conflict of Interest

The authors have no conflicts of interest to disclose.

## Notes

### Competing Interest Statement

The authors have declared no competing interest.

